# Evolution of intrinsic transcriptional terminators in their genomic context

**DOI:** 10.1101/2024.07.17.603904

**Authors:** David Toledo-Aparicio, Stepan Denisov, Charles Bernard, Rodrigo AF Redondo, Caroline C Carter, Zahra Khomarbaghi, Dorothea A Pittrich, Anna Nagy-Staron, Calin C Guet, Mato Lagator

## Abstract

Transcriptional gene expression relies on two fundamental processes – transcription initiation and termination. Transcriptional termination is essential for the coordinated control of gene expression. Intrinsic termination, the major mechanism of transcriptional termination in bacteria, relies on the sequence-dependent folding of nascent mRNA into a hairpin structure that enables RNA polymerase to dissociate from DNA. Despite their central role in regulating expression in prokaryotic genomes, the conservation of intrinsic terminators and the factors that shape their evolution remain poorly understood. Here, we combine comparative genomics with experimental measurements of terminator function, to study the conservation of intrinsic terminators between *Escherichia coli* and *Salmonella enterica* subsp*. pullorum*. While the two species are closely related, the sequence and function of most terminators are not conserved, with less than 20% of all terminators having an identifiable ortholog – in stark contrast to 60% for coding sequences. Terminators with higher sequence conservation also had more conserved function, indicative of stabilizing selection. The local genomic context shapes the evolution of intrinsic terminators, as their sequence conservation is dependent on the conservation of the upstream gene, while their function is affected by the distance to the downstream gene. Ultimately, any theory of gene regulatory network evolution ought to account for how transcriptional terminators evolve.

## Introduction

Regulation of gene expression is an essential feature of all living organisms, enabling them to respond to their environment and coordinate internal cellular processes ^1^. In prokaryotes, a key aspect of gene regulation involves terminating transcription at specific locations around the genome, which enables operon-specific control of gene expression levels by isolating operons from the regulatory context of surrounding genes and promoters ^2^. Transcriptional termination is achieved through two mechanisms ^3^: (i) rho-dependent termination, which requires the binding of the Rho protein to nascent mRNA; and/or (ii) rho-independent (intrinsic) termination, which relies on the passive formation of a secondary RNA structure that induces the disassociation of RNA polymerase from DNA.

The molecular mechanisms of transcriptional termination have been extensively studied almost exclusively *in vitro;* and hence outside of their cellular context ^3^. While the question of how termination is achieved at a mechanistic level has been studied and understood in great detail, the relationship between sequence and the *quantitative* function of a terminator – *i.e*., how efficiently it terminates transcription – has received much less attention ^4–6^. Unsurprisingly, not all terminators have the same function, as some are more efficient than others in their ability to terminate transcription ^4^. Such differences in terminator efficiency introduce another level of complexity to bacterial gene regulation, whereby less efficient termination enables transcriptional read-through of downstream genes ^7,8^. Transcriptional read-through from even efficient terminators can alter the logic of a gene regulatory network ^9^ or even affect the fitness of an organism ^8^. Similarly, mutations in *rho* can be an important force enabling the evolution of downstream promoters ^10^, suggesting that terminators could be selected to have specific efficiency. For example, terminators upstream of operons that require tight control of their expression could be selected to be very efficient. Conversely, it is possible that some terminators would be selected to have lower efficiency if the transcriptional read-through of downstream operons has a beneficial impact on fitness.

What are the factors that can shape selection on terminator efficiency? On the one hand, selection for terminator efficiency ought to depend on the specific operon(s) surrounding the terminator and how tightly those operons need to be regulated. If dominant, this source of selection on terminator efficiency would have to be studied in an operon-specific manner. On the other hand, a set of generalizable factors that depend on the genomic context of the terminator could shape the selection for termination efficiency. The immediate genomic context of a terminator includes, among other factors, the orientation of the genes around it and the distance between the terminator and those genes. How the genomic context shapes terminator evolution remains poorly understood, in part because of the scarcity of studies investigating the evolutionary history of terminators ^11,12^. In this study, we set out to investigate the relationship between the genomic context of the terminator, its efficiency (function), and the evolution of its sequence. We focus on *rho*-independent termination, also referred to as intrinsic termination, because the relationship between sequence and RNA structure is well understood in this system ^13,14^, allowing computational predictions of intrinsic terminators (ITs) across the genome ^15^.

## Results

### Uncovering structural properties and genomic context of ITs

The study of IT evolution requires annotating ITs directly from genomic sequence, obtaining their structural information, and then identifying their genomic context. To automatize this process, we developed an integrated pipeline – ITMiner. To identify IT sequences in a bacterial genome, ITMiner starts with an *in silico* approach, RNIE ^15^, which identified 1665 ITs in the *Escherichia coli* MG1655 genome. For ITMiner, we adapted RNIE to generate a conservative list of ITs by minimizing the number of false positive predictions by: (i) considering ITs identified on opposite DNA strands, which have >90% overlap in sequence and contain an identifiable polyuridine (poly-U) tract on their respective strands, to be one IT capable of acting in both directions instead of two independent ITs. Note that the orientation of the IT (*i.e.,* which strand the terminator is acting on) is determined by its poly-U tract, meaning that some ITs can be bidirectional if they have a poly-U tract on both strands; (ii) removing all ITs predicted inside coding sequences (for more details, see Materials and Methods). These conservative criteria reduced the list of ITs in *E. coli* to 644 (Supplementary Data 1), which we then compared to two lists of experimentally derived ITs. Our approach predicted 72% (448/623) of ITs identified by Conway *et al*. ^16^ and 61% (163/267) of ITs found in RegulonDB ^17^, as well as 154 ITs that were not identified by either of these datasets. ITMiner then identifies the structural features of each IT by calculating the most stable 2D folding structure using the Vienna Package ^14,18^. Based on this structure, ITMiner obtains the sequence of the stem, loop, and poly-U tract. Finally, ITMiner provides the position of each IT in the reference genome.

First, we characterized the properties of the 644 ITs identified in *E. coli*. Each IT consists of a stem, loop, and a poly-U tract (Fig.1A), and we described the distribution of their lengths and GC contents (Fig.1B–E). The exact length of the poly-U tract as identified by RNIE ^15^ is not precisely defined, which is why we did not analyze the GC content and length of poly-U tracts. In contrast to the “model” ITs used to study the molecular mechanisms of IT function ^3,19^, *E. coli* ITs tolerate a wide diversity of GC contents in loops and stems. GC content in stems has a wide distribution that appears log-normal, even though stems are traditionally assumed to be GC-rich ^2,13^.

**Figure 1.**
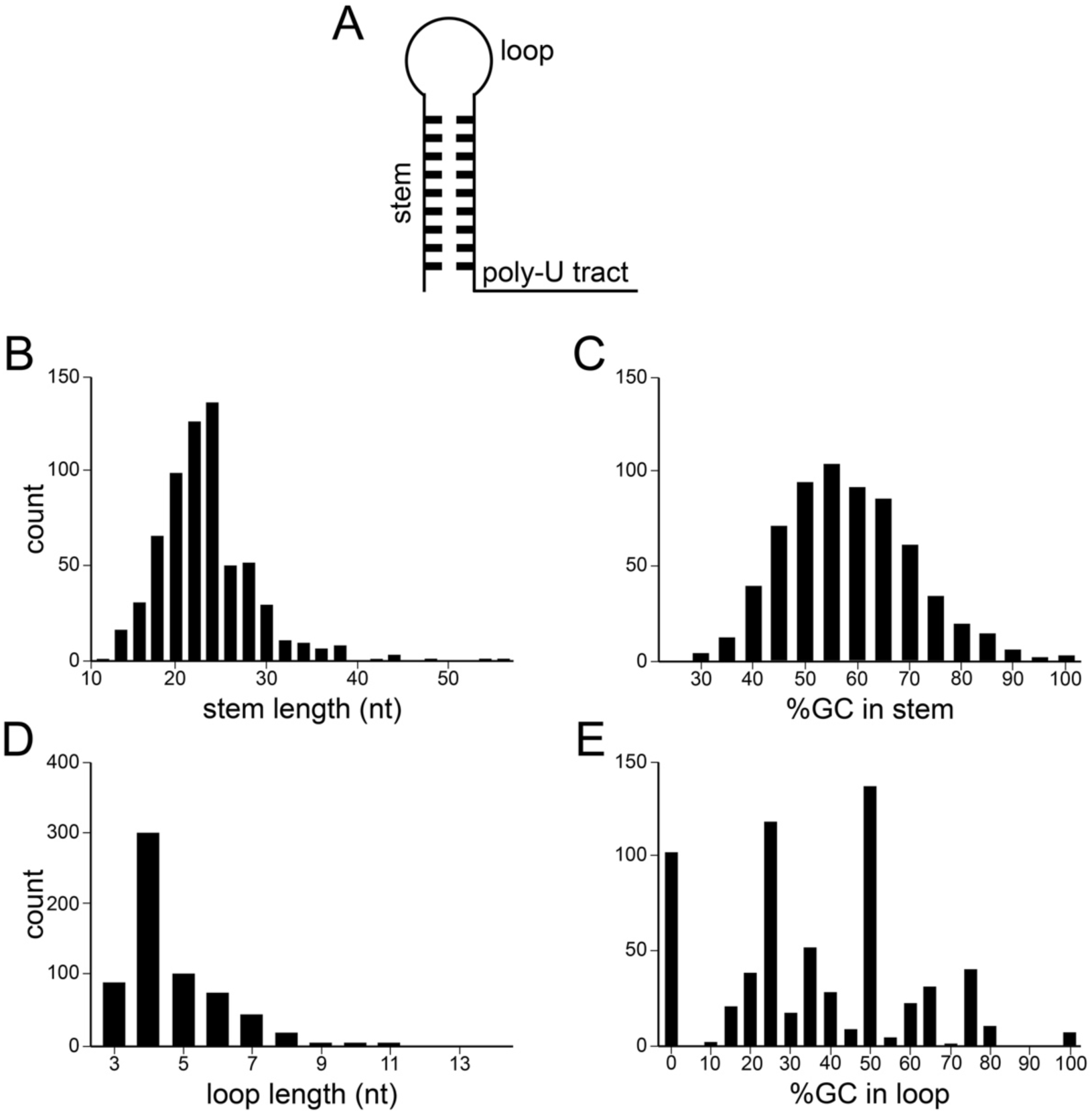
Intrinsic terminators (ITs) in the *E. coli* genome. (**A**) General structure of ITs, consisting of stem, loop (which together form the hairpin), and polyuridine (poly-U) tract. These are the key features that the RNIE algorithm ^15^ uses to identify ITs in a genome. Distributions of stem lengths (**B**) and their GC content (**C**), as well as loop lengths (**D**) and their GC content (**E**) are shown. Length and GC content distributions for the poly-U tract are not shown because, mechanistically, it is not well understood what defines the boundaries of poly-U tracts. Thus, their size, as determined by RNIE, is an estimate.

Based on the output from ITMiner, we characterized the genomic context of ITs in *E. coli* according to: (i) the orientation of the genes immediately upstream and downstream of an IT, meaning that the orientation can be “head-to-tail”, “head-to-head” or “tail-to-tail” (Fig.2A); and (ii) the distance between the IT and the genes immediately upstream and downstream of it (Fig.2B). The fact that we identified only three tail-to-tail ITs, a genomic context where the existence of an IT is not expected, serves as further validation of the robustness of the ITMiner pipeline used to predict ITs directly from sequence. The distance between an IT and its immediate upstream gene is significantly shorter than the distance to the immediate downstream gene (Fig.2B; t_1286_ = 175, P < 0.0001), suggesting that the IT location relative to its genomic context might be under selection. We identified twelve intergenic regions that contained two consecutive non-overlapping ITs on the same strand, potentially suggesting stronger selection for the transcript to be terminated in that region.

**Figure 2.**
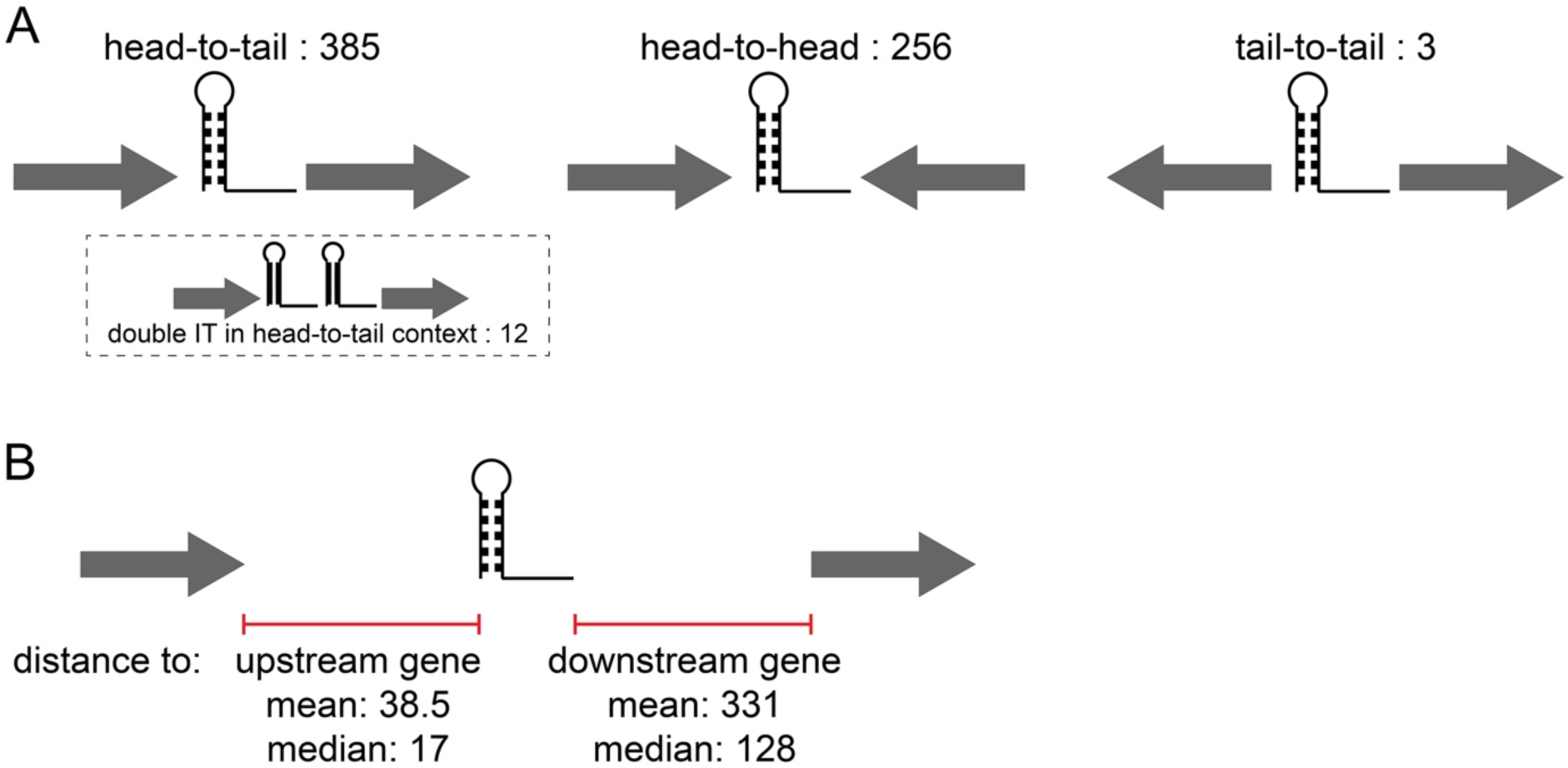
Genomic context of intrinsic terminators in *E. coli*. Two factors define the genomic context of an IT. (**A**) The orientation of genes immediately surrounding the IT, which can be head-to-tail, head-to-head, or tail-to-tail. We identified twelve intergenic regions that contained two adjacent ITs, all of which were head-to-tail. The IT count in each category (total: 644) is shown. (**B**) Distance from the IT to the genes immediately upstream and downstream, irrespective of their orientation (here shown as head-to-tail for illustration purposes). Bidirectional ITs (those that act on both strands) were not included in this analysis as there was no difference between upstream and downstream genes in that case. Arrows represent genes, and stem/loop/polyuridine tract structure the IT.

### Efficiency of E. coli ITs

To explore the functional diversity of ITs, we constructed a plasmid system that allowed us to measure the efficiency of transcriptional termination for individual ITs in a cellular context. The system consisted of a strong, inducible promoter that controlled the expression of a cyan fluorescent protein (*cfp*) gene, a terminator sequence isolated by RNaseE sites, and a yellow fluorescent protein (*yfp*) gene (Fig.3A). The system was placed in a low-copy number SC101* plasmid ^20^. We selected 20% of *E. coli* ITs (129 from the list of 644 ITs identified computationally), inserted each into the plasmid, and calculated their efficiency by measuring how they modulated YFP expression levels. None of the chosen ITs were bidirectional but were otherwise selected at random.

**Figure 3.**
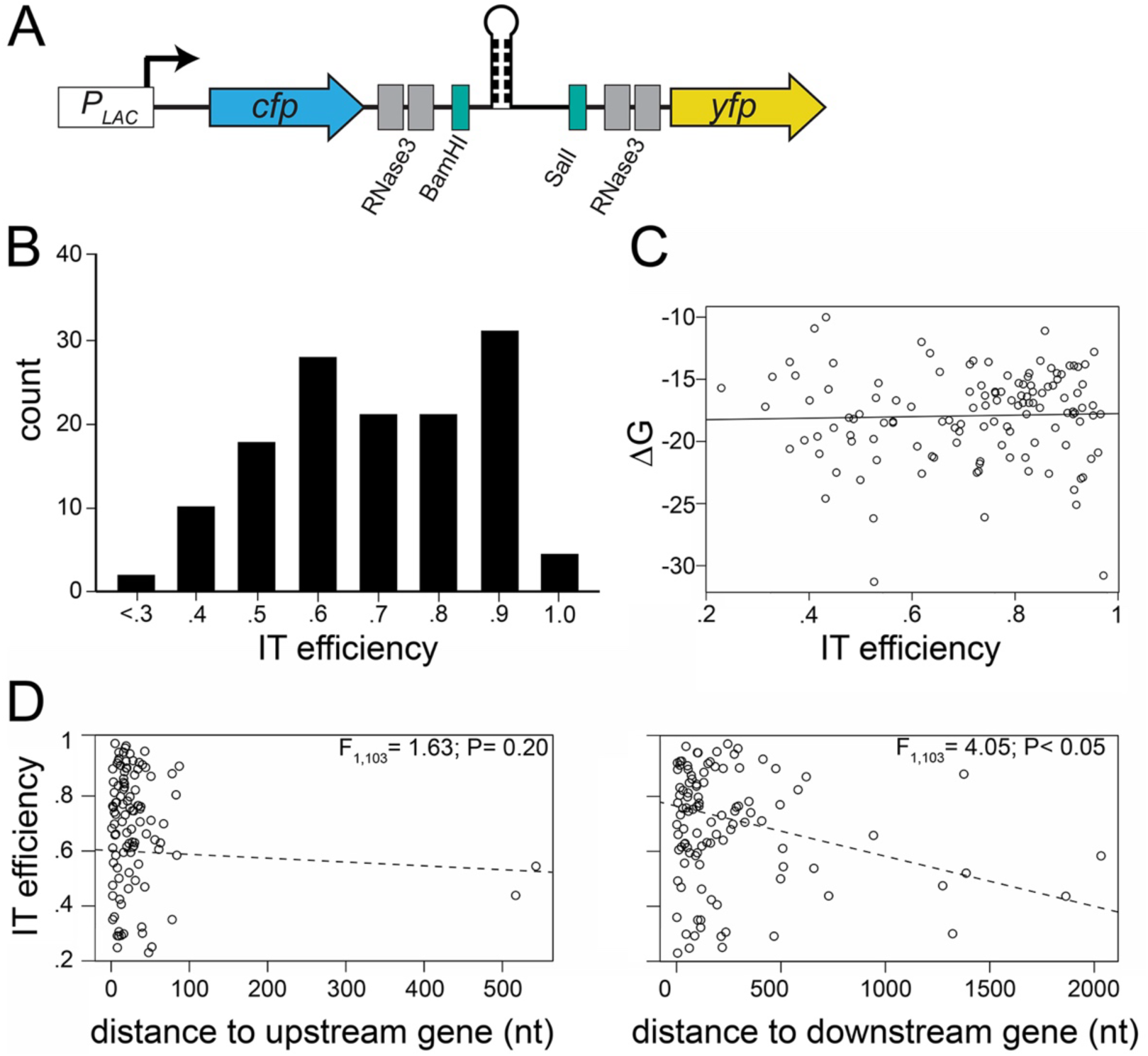
Efficiency of *E. coli* intrinsic terminators (ITs). (**A**) Genetic cassette designed to measure IT efficiency, consisting of *cfp* and *yfp*, separated by RNase3 and unique restriction sites (BamHI and SalI), under the control of an inducible *P_LAC_* promoter on a low-copy plasmid. This system allows the insertion of any desired IT and determination of its efficiency by comparing the extent to which it reduces relative YFP expression compared to a control lacking any IT. RNase3 sites separate *cfp* and *yfp* transcripts, ensuring that differences in fluorescence are not due to an IT introducing any unique post-transcriptional modifications or changing mRNA stability. (**B**) Distribution of IT efficiencies across 129 measured *Escherichia coli* ITs. (**C**) Correlation between the Gibbs free folding energy (Δ*G*) of ITs and their efficiency in *E. coli* (F_1,103_ = 0.12; P= 0.73; R^2^ = 0.001). (**D**) Correlation between IT efficiency and the distance to either the closest upstream (left plot) or downstream (right plot) gene. Circles represent the mean efficiency estimated from three independent replicate measurements. The dotted line represents the linear regression. The correlation between distance to upstream gene and IT efficiency (left plot) was also not significant by excluding the two outliers (which were both in the head-to-tail orientation).

The tested ITs exhibited a wide range of efficiencies (Fig.3B). Their efficiency did not correlate with the Gibbs free folding energy, meaning that the efficiency of ITs is not solely determined by their structural stability (Fig.3C). 29 ITs were so inefficient that we could not statistically separate them from a control sequence that did not contain an IT, possibly due to our use of fluorescence measurements at population level or a false positive prediction of ITs in our pipeline. Consequently, they were excluded from further analyses. These findings suggest that selection for high IT efficiency is not acting on all ITs. Rather, some ITs are likely not under strong selection to be highly efficient or might be selected specifically to have intermediate or low efficiency.

We asked whether IT efficiency depended on their genomic context. We found that the orientation of the genes surrounding the IT did not significantly impact their efficiency (comparison carried out between head-to-tail and head-to-head ITs, as our dataset included only one tail-to-tail IT: t_102_ = 0.65, P = 0.51). Then, we examined if the distance from an IT to its immediate upstream or downstream gene correlated with efficiency, excluding bidirectional ITs from this analysis. We found a weakly significant, negative correlation between IT efficiency and the distance to downstream genes (F_1,103_ = 4.05, P < 0.05) (Fig.3D), meaning that as the distance between ITs and downstream genes increased, IT efficiency decreased. This implies that the selection for IT efficiency (its function) might be driven by the requirement to transcriptionally isolate the downstream gene, which can be achieved either by selecting for a more efficient IT (Fig.3D) and/or by increasing the distance between operons, which leads to an increased probability of spontaneous RNA polymerase dissociation and hence termination (Fig.2B).

### Conservation of ITs between two closely related species

As opposed to promoters, which determine gene expression levels and whose evolutionary history has been studied in more detail ^21–23^, how selection acts on ITs remains a black box. In other words, it is unknown whether selection acting on IT sequences is strong, weak, or neutral. Several challenges exist when trying to understand the history of selection acting on ITs. First, their sequences are very short, typically 25–50 nucleotides, posing a difficulty when constructing alignments and inferring phylogeny because the tools for alignment and phylogenetic inference typically require longer sequences to reliably infer the number of substitutions and, hence, relatedness between two sequences. Second, in contrast to protein sequences, the function of an IT is inherently linked to its genomic context, as the function of an IT is to impact transcription of downstream genes. In other words, the function of an IT is inherently connected to the genes in its immediate context, the distance to them, and their orientation. This means that their evolutionary history is particularly sensitive to genomic rearrangements and horizontal gene transfer, further complicating the application of traditional phylogenetic approaches. Finally, many common measures of selection rely on identifying whether a specific mutation is synonymous or not. This is easily defined for proteins, where the redundancy in the genetic code provides a clear delineation between synonymous and non-synonymous mutations (although, this metric ignores the possible selection for codon usage ^24^). However, despite some attempts ^4^, the relationship between IT sequence and function is poorly understood, making it very difficult to estimate whether a mutation in an IT is neutral ^11^.

Thus, to avoid these difficulties, we adopted a more qualitative approach based around global measures of IT conservation between two closely related species, *Escherichia coli* and *Salmonella enterica subsp. pullorum* (from now on, *S. pullorum*) (for the somewhat odd choice of the second species, see Materials and Methods).

### Conservation of ITs based on sequence homology

First, we aimed to identify homologous IT pairs between *E. coli* and *S. pullorum*. We used BLAST on each *E. coli* and each *S. pullorum* IT sequence from our computationally derived list of ITs (Supplementary Data 1 and 2). We considered an *E. coli* IT to have a homolog in the *S. pullorum* genome when we found a significant BLAST hit in the *S. pullorum* genome (note that BLAST was conducted against the entire genome of the opposing species not just against the list of identified ITs). Then, an analogous definition was applied to identify whether each *S. pullorum* IT had a homolog in the *E. coli* genome. Note that homology does not need to be reciprocal, as an *E. coli* IT sequence can have more than one significant BLAST hit in the *S. pullorum* genome. We considered *E. coli* and *S. pullorum* ITs to be orthologous (meaning that they likely evolved through a speciation event) when they were the top BLAST hit of each other (a.k.a. bidirectional best hit approach ^25^).

For almost 85% of *E. coli* and 80% of *S. pullorum* ITs we could not identify an ortholog (with Jaccard indexes in both genomes > 0.4, Fig.4A), suggesting the general absence of stabilizing selection acting on IT sequences during the evolutionary history of these two species. For comparison, only 37% of *E. coli* and 33% of *S. pullorum* protein-coding genes do not have an ortholog in the other species (with Jaccard indexes in both genomes > 0.4). Orthologous IT pairs had a varied number of mismatches, with most being either identical or having 1–3 mismatches (Fig.4A). We did not identify any patterns in the location of mismatches, as they were evenly spread across stems, loops, and poly-U tracts.

**Figure 4.**
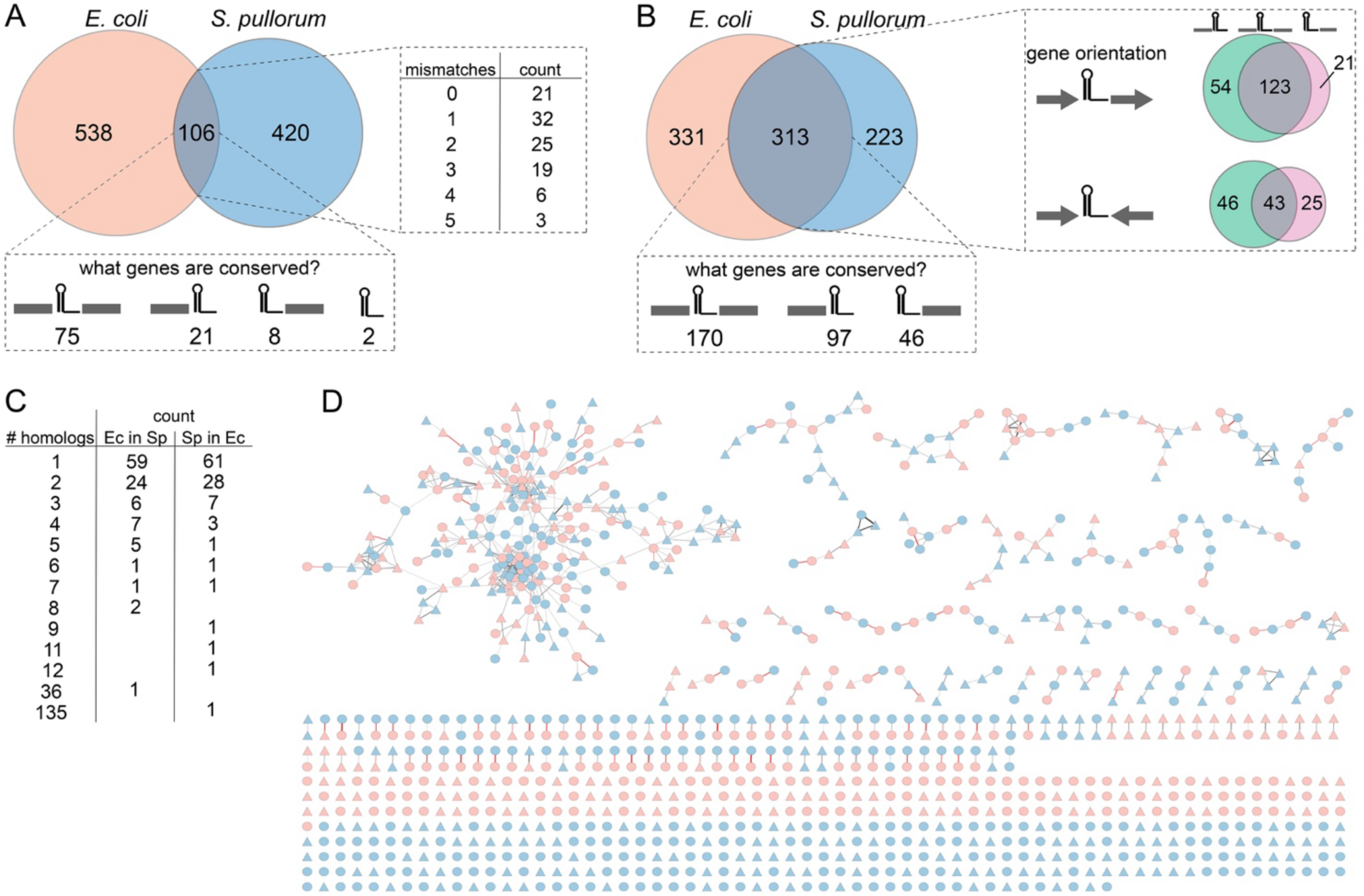
Conservation of intrinsic terminators (ITs) between *E. coli* and *S. pullorum*. (**A**) Venn diagram showing how many ITs had an identifiable ortholog. An IT in *E. coli* had an ortholog in *S. pullorum* when they were each other’s top BLAST hits. The table shows the number of mismatches between all identified ITs. The box below shows information on whether the ITs had an orthologous upstream gene, downstream gene, both, or none. Gray rectangles represent the location (upstream or downstream) of conserved genes with respect to the IT. **(B**) Venn diagram showing ITs present at the same genomic location in both species – *i.e*., we identified an IT in both species immediately upstream or downstream (or both) of a conserved gene. Additional Venn diagrams show how the number of ITs that had either upstream, downstream or both genes conserved depended on the orientation of those genes. (**C**) Number of homologs identified for each IT from *E. coli* (Ec) in the *S. pullorum* (Sp) genome and *vice versa*. (**D**) Sequence Similarity Network (SSN) of the 664 *E. coli* and 516 *S. pullorum* ITs. Nodes correspond to ITs and edges depict high sequence similarity links between two ITs. ITs are colored according to their species origin (blue for *E. coli*; red for *S. pullorum*), shape indicates the genomic context (circle for head-to-tail, triangle for head-to-head). The width of edges are proportional to the −log10(E-value) of the alignment between two ITs, with red edges corresponding to BBHs (*i.e.*orthologs) between *E. coli* and *S. enterica* identified through the BLAST approach.

Remarkably, approximately 6% of *E. coli* and 4% of *S. pullorum* ITs had more than one homolog with E value < 0.0005 (Fig.4C). One *E. coli* IT (Cha381) had 36 homologs in the *S. pullorum* genome, while one *S. pullorum* IT (Nur16) had 135 homologs in the *E. coli* genome. The existence of more than one homolog between *E. coli* and *S. pullorum* could be due to rapid and widespread horizontal transfer and/or duplication events. Surprisingly, the corresponding coding regions in the immediate context of these ITs did not exhibit such one-to-many mappings.

To further investigate the relatedness between ITs of the two species, we conducted an all-against-all alignment between the 1,180 ITs in our dataset (644 *E. coli* and 536 *S. pullorum* ITs) using the exact Smith-Waterman algorithm. This approach gave us more precise information about sequence similarity between ITs but, in contrast to the BLAST approach, was constrained only to the ITs identified by ITMiner instead of the entire genome. In line with the BLAST-based analysis (Fig.4C), the all-against-all approach identified most ITs (525) as orphans or having only one highly similar sequence (338) (Fig.4D). We identified 26 clusters that had more than 10 highly similar ITs, including a very large cluster that included both Cha381 and Nur16. Interestingly, ITs from one species were as likely to be related to ITs of the other as to those from the same species (assortativity coefficient of 0.18). Head-to-tail ITs were not preferably connected to other head-to-tail ITs (assortativity coefficient of 0.05), implying that ITs are regularly repurposed in a different genomic context in the process of their propagation and diversification.

We wanted to better understand the evolutionary history of the largest connected group of ITs (Fig.4D), which included the two ITs with the highest number of homologs (Fig.4C). We considered the phylogenetic relatedness between all the BLAST hits (homologs) for these ITs, both in *E. coli* and in *S. pullorum*, and asked whether the two IT homologs were related to each other, and, if so, whether the species split predated any duplication or sequence diversification events. All Nur16 and Cha381 homologs were indeed related to each other. Remarkably, sequences closer to Cha381 clustered together, as did the sequences closer to Nur16, irrespective of the species the ITs were found in (Fig.5). This suggests that the two ITs diversified prior to the *Escherichia* and *Salmonella* split. In the much better annotated *E. coli* genome, all BLAST hits of the two ITs overlapped in sequence with an insertion sequence *IS*186 ^26,27^. The close association between this group of highly similar ITs and *IS*186 likely enabled their rapid and large-scale intra-genome propagation.

**Figure 5.**
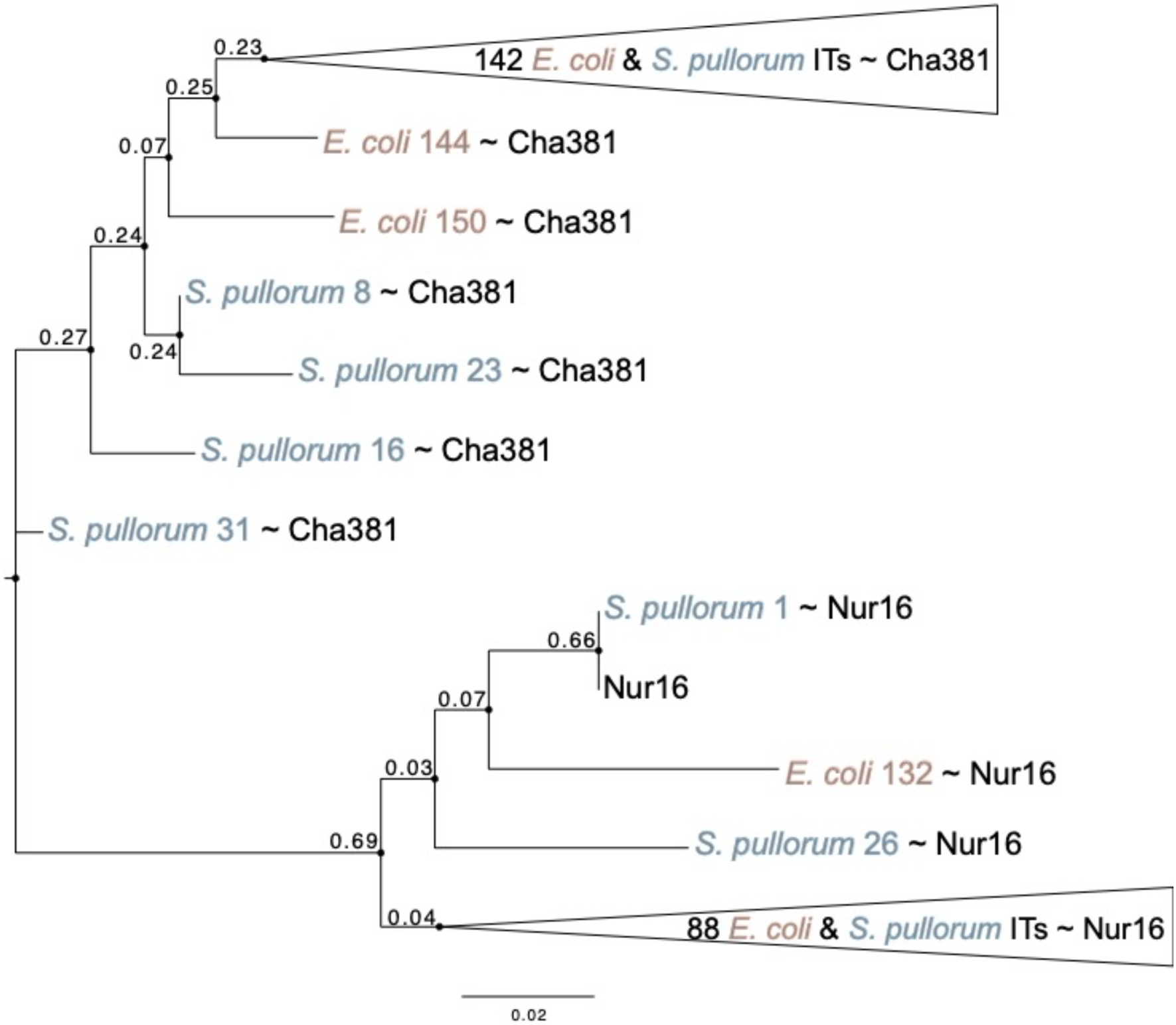
Phylogeny of two intrinsic terminators (ITs). We constructed an alignment of all BLAST hits of the two ITs with highest number of homologs, in both *E. coli* and *S. pullorum* and used it to construct a neighbor-joining phylogenetic tree to represent the relatedness sequences. We only show branches with bootstrap values > 0.1, as many sequences were identical or differed in only one point mutation. The species in which an IT was found is contained in its name and shown in orange for *E. coli* and blue for *S. pullorum*. Numbers in orange or blue correspond to the name of the IT in Supplementary Data 1 and 2. Whether an IT was closer in sequence to Cha381 or to Nur16 is indicated by tilde (∼).

### Conservation of ITs based on genomic context

To further probe the role of the local genomic context on IT evolution, we used a second measure of IT conservation based on the genes that surround an IT in *E. coli*, asking if we could identify an IT within the same genomic context in *S. pullorum*. We found that 170 ITs were present in the same context in *E. coli* and *S. pullorum*, meaning they had orthologous upstream and/or downstream genes even though the ITs themselves were not necessarily homologous to each other. Of these, 97 contained only an orthologous upstream gene, while 46 contained only an orthologous downstream gene (Fig.4B). In other words, an IT is least likely to be found in the same genomic context if associated only with the downstream gene. A possible explanation is that, if changes in the function or the expression of genes between *E. coli* and *S. pullorum* are common, then the link between downstream gene and IT efficiency (Fig.3D) could be driving more rapid events of positive selection and hence lower IT conservation associated with downstream genes (Fig.4C). Alternatively, it could be due to the longer distance between an IT and its downstream gene increases the likelihood of them being decoupled due to a rearrangement event. Interestingly, not all 106 homologous ITs had both upstream and downstream genes conserved (Fig.4A). In fact, 21 and 8 homologous ITs had only the upstream and downstream gene conserved, respectively. Furthermore, two ITs identified as identical homologues were not found within the same genomic context, meaning that neither their upstream nor downstream genes were homologous between *E. coli* and *S. pullorum*, suggesting the possibility of a short-sequence genomic re-arrangement ^28^. Finally, we found that head-to-tail ITs are more likely to preserve their genomic context, compared to head-to-head ITs (Fig.4C; *χ*_2_ = 82, P < 0.0001) – suggesting that transcriptionally insulating genes is more important when genes are in the head-to-tail context, *i.e.,* found on the same DNA strand.

Next, we asked if a conserved genetic context between *E. coli* and *S. pullorum* – namely whether the upstream, downstream, or both genes in one species had an ortholog in the other in the same genomic configuration relative to the IT – affected the probability of finding an IT ortholog in that context. In other words, we explored the extent to which the genomic context surrounding an IT was conserved depending on the conservation of the IT itself. Unsurprisingly, conserved ITs were more likely to also be found in a conserved context, likely due to selection acting on the whole architecture (IT and its associated genes) (Table 1). The majority of conserved ITs had both the upstream and downstream genes conserved as well. Less frequently only the upstream gene was conserved, and least frequently only the downstream gene – likely due to the greater average distance between ITs and downstream genes (Fig.2B), facilitating genomic rearrangements. This was also true for non-conserved ITs (*i.e.,* when a non-homologous IT was found in the same genomic context). Interestingly, these trends were only observed in head-to-tail context, as ITs were rarely conserved when found in head-to-head context, even if that context was itself conserved.

What could be the biological role of this finding? The head-to-tail orientation leads to the upstream gene having a degree of transcriptional read-through into the downstream gene and thus, a degree of regulatory connection. Therefore, our findings reaffirm the importance of transcriptional read-through as a force shaping gene regulatory network evolution and function ^9,29^, while individual promoter evolution can be more local ^30,31^. In contrast, ITs in head-to-head context experience weaker selection pressure, likely due to readthrough having a smaller effect on the expression levels of a gene on the opposite strand.

These results indicate that the global factors arising from the genomic context did not result in strong stabilizing selection on IT sequences, as only ∼20% of *E. coli* ITs had at least one ortholog in the closely related species *S. pullorum*. The hypothesis of weak stabilizing selection acting on IT sequences is further supported by the finding that 50% of ITs are found within the same genomic context but many of them are not homologous (Fig.4), suggesting a high substitution rate within ITs and/or intensive genomic rearrangements/horizontal transfers. Our findings suggest that stabilizing selection acts on ITs predominantly within a head-to-tail context and when both upstream and downstream genes are conserved, while ITs evolve more rapidly when found in other genomic contexts.

### Conservation of IT function

Finally, we wanted to study how the function of ITs – their efficiency – is conserved between two closely related species, *E. coli* and *S. pullorum*, and how factors associated with the genomic context might alter IT efficiency. To address this question, we randomly selected 45 ITs of *S. pullorum* that had an identifiable (but not identical) ortholog in *E. coli*, inserted them into our plasmid system and measured their efficiency. The distribution of mismatches among these 45 ITs mirrored the distribution of mismatches observed among all ITs (compare Fig.6A and Fig.4A). We measured IT efficiency in both hosts and found that terminator efficiency was not host-specific (Fig.6B) – an expected finding given the high sequence homology between all genes involved in transcription and termination (Supplementary Table 1). For this reason, the ensuing analyses were conducted based on efficiency measurements in *E. coli*.

**Figure 6.**
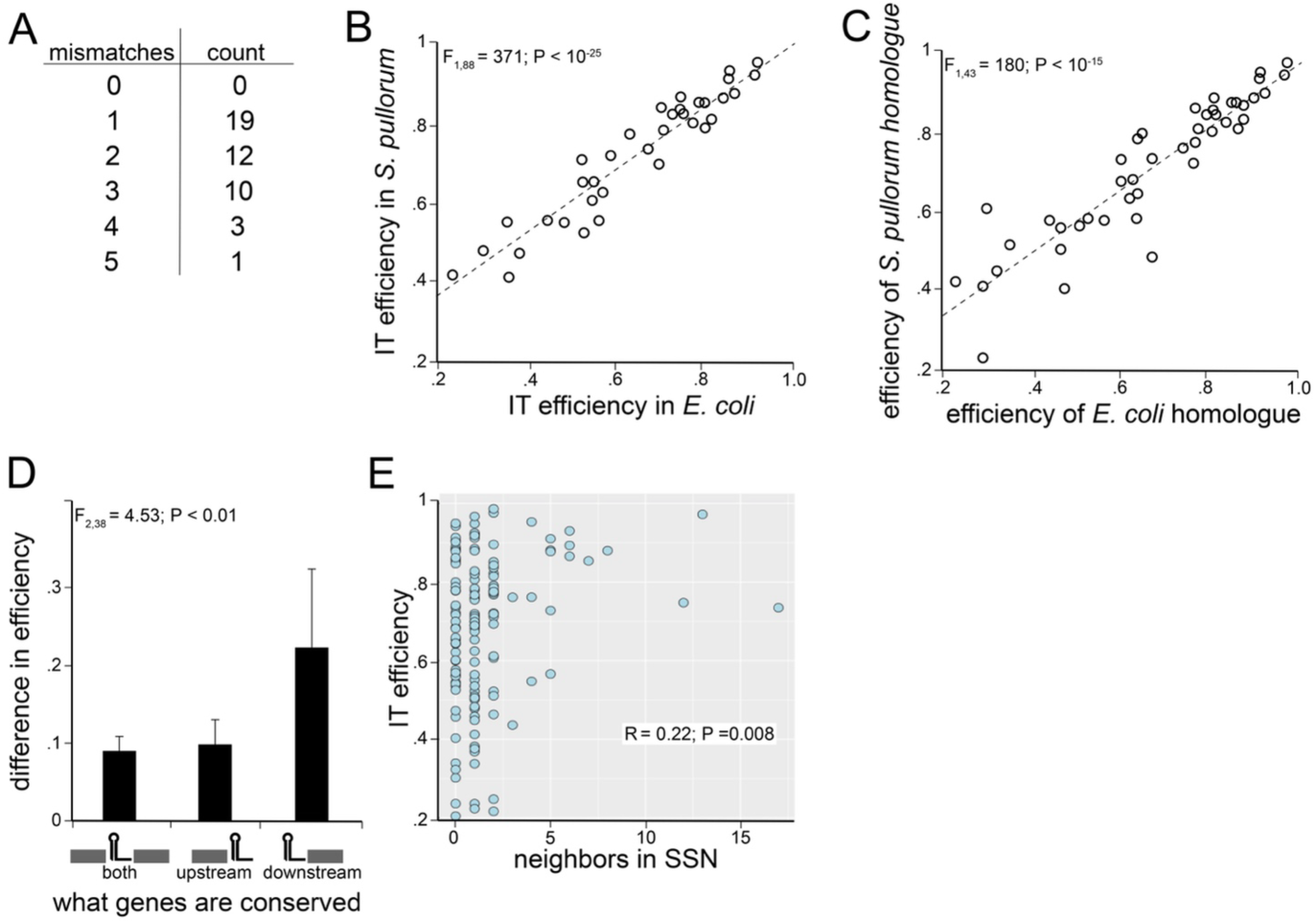
Efficiency of homologous intrinsic terminators (ITs). (**A**) Distribution of mismatches between 45 homologous IT pairs, for which termination efficiency was measured. (**B**) Correlation between IT efficiency of all 90 ITs (45 homologous pairs) measured in *E. coli* and in *S. pullorum*. Black circles represent the mean efficiency from three replicate measurements in both hosts, error bars show the standard deviation. The dotted line represents the linear regression. (**C**) Correlation between efficiencies of homologous IT pairs. Each circle is the mean of three replicate measurements in *E. coli*. The dotted line represents the linear regression. (**D**) Absolute difference in mean efficiency between each homologous IT pair, as a function of whether upstream, downstream, or both genes were conserved. Black bars represent mean absolute differences in efficiency, error bars show the standard deviation. Anova test results are shown. (**E**) Correlation between the number of neighbors an IT has in the SSN (Fig.4D) and its efficiency.

Most orthologous ITs exhibited similar efficiencies (Fig.6C). The differences in IT efficiency between orthologs did not significantly depend on the number of mismatches between them (F_4,33_ = 2.00, P = 0.12). However, even though the differences in IT efficiencies were generally small, they were sufficient to investigate whether genetic context impacted IT efficiency. The difference in the efficiency between orthologous ITs did not depend on the orientation of genes surrounding the IT (t_44_ = 0.14, P = 0.89; compared between head-to-tail and head-to-head ITs only). However, it was significantly affected by the conservation of genes surrounding the IT (Fig.6D). Specifically, having either only the upstream gene or both the upstream and downstream genes conserved, resulted in significantly smaller differences in efficiencies between homologs compared to when only their downstream gene was conserved (F_2, 38_ = 4.53, P < 0.01). Interestingly, we observed a correlation between the number of relatives an IT has (*i.e.,* the number of SSN neighbors, Fig.4D) and its efficiency (Fig.6E), suggesting that higher IT efficiency impacts the likelihood of an IT propagating in the genome and/or the likelihood of it being conserved.

## Discussion

We set out to identify the genomic factors that shape IT efficiency and the conservation of IT sequence and function between two closely related bacterial species. Through a combination of qualitative, functional, and comparative genomics analyses, we unraveled some of the complex factors that shape IT evolution and drive rapid diversification of IT sequences, even between two closely related species. Most *E. coli* ITs did not have a clearly identifiable homolog in *S. pullorum*, while those ITs that did, had a strongly preserved function (efficiency). We found that the conservation of the upstream gene and the distance between the IT and the downstream gene play a key role in preserving IT sequence and function.

The complex set of factors that affect the evolution of ITs point to a key finding: their sequence and, implicitly, their function, change rapidly. This is best observed in the fact that, even though we chose two closely related species, less than 20% of all ITs had an ortholog – in stark contrast to ∼65% for coding sequences. It is possible that ITs have a high rate of sequence variation because most are under very weak selection or evolve neutrally. If true, this would suggest that the effect of most ITs on transcriptional gene expression levels and transcriptional read-through does not impact organismal fitness. The wide diversity of experimentally measured IT efficiencies, which mirrored those from a previous study ^4^ (Fig.3B), coupled with low sequence conservation, provides further indirect evidence that altering IT efficiency might not always impact organismal fitness.

Alternatively, ITs could be under strong directional selection, whereby the inability to identify homologs for most ITs stems from strong selection pressure to alter their function acting in at least one of the lineages we studied. Notably, the ∼20% of *E. coli* ITs that did have an identifiable sequence ortholog also had conserved function (Fig.6C), suggesting that stabilizing selection might be acting on at least some IT sequences.

One clear message that emerges from these results is that phylogenetic methods capable of dealing with variations in short sequences within a corresponding context need to be developed in order to better discern the sources of selection that act on IT sequences. These methods need to also be able to account for the qualitatively different effects of mutations on ITs, compared to proteins. Studies of protein evolution critically rely on differentiating between synonymous mutations and those resulting in amino acid changes. Even such broad sequence-to-function mapping is lacking for ITs, which hampers the study of their evolution.

A key factor shaping the function and sequence preservation of ITs is the identity and function of the genes/operons that are immediately upstream and downstream ^7,11^. Our study focused on broader and more generalizable factors of ITs’ genomic context and hence did not explore any gene-specific effects. It is plausible that gene-specific effects are the major drivers of IT evolution, potentially explaining why so few ITs have a direct sequence homolog between two closely related species. This is particularly true given the high rate of genomic re-arrangements and shuffling commonly observed in bacteria ^28,32^, which could be providing new genetic context upstream and/or downstream of a given IT.

A potential problem with our approach is that we use bioinformatics to predict ITs across genomes ^15^. While this allowed us to explore IT conservation between two species, one of which is not a well-annotated model organism, it also means that our IT library may be incomplete and/or contain false positives. In fact, some of the ITs whose function we experimentally measured showed no significant levels of transcriptional termination, at least based on the population-level measurements of fluorescence adopted in this study. While this could be because of the reduced sensitivity of our experimental assay, it could also be indicative of the false positive rate in the algorithm used to identify ITs. Overcoming this difficulty is not trivial, as there is no universal experiment that would identify all ITs, because detecting an IT requires its upstream promoter to be active – a condition that cannot be easily achieved experimentally for each promoter in a genome ^16,33^.

The evolutionary history of ITs remains poorly understood, with existing studies focusing either on broad approaches that analyze only the number and structure of ITs across bacterial phylogeny ^12^, or those that focus on the evolutionary history of individual ITs ^11^. What these studies lack is a reference to the actual function of individual ITs – their efficiency. Combining a sequence conservation-based approach with measurements of IT efficiency allowed us to identify genomic context-dependent factors that shape IT evolution. The development of better phylogenetic techniques to study evolutionary dynamics of short sequences, combined with better tools for identifying ITs and high-throughput functional approaches to measure IT efficiency, will be key in unravelling the mystery of IT evolution. Early work in synthetic biology ^29^ suggested that the context of ITs can be essential in determining the function of genetic networks ^9^. Without unravelling the evolutionary puzzle of ITs, we cannot achieve full understanding of how gene regulatory networks evolve nor precisely engineer the genetics of cells.

## Materials and methods

### Obtaining IT sequences from genomes

To obtain IT sequences from bacterial genomic sequences, as well as relevant contextual and structural information about each IT, we developed an integrated pipeline, ITMiner (code for ITMiner can be found in Supplementary Data 3). First, ITMiner utilizes the RNIE algorithm ^15^ to identify potential IT sequences in the genome. The algorithm has two modes – “gene” and “genome” – which it uses to scan the genome for potential IT sequences, as described by Gardner *et al.* ^15^. We used both modes and combined their outputs to identify 1665 candidate sequences in the *E. coli* K12 MG1655 genome (NC_000913). This initial list contained numerous sequences that were identified as two ITs, containing identical stem and loop but with poly-U tracts of different lengths. This occurred because the structure of the poly-U tract is not as clearly defined as that of the stem, allowing sequences with varied lengths to be identified as potential poly-U tracts. In such instances, we selected only the longest of the identified overlapping sequences, reducing the number of candidates to 1065. Then, we removed all candidate ITs found within coding regions, bringing the number to 694. While some predicted ITs within coding regions might be functional, we opted for a more conservative approach and excluded them as, generally, transcription should not be terminated in the middle of a gene. Finally, a number of candidate ITs were overlapping in sequence, but were identified on opposite strands. This is because a sequence that forms a hairpin on one strand will by definition also form one on the other; thus, the only distinguishing feature bioinformatically will be the presence or absence of the poly-U tract. We identified the structural features of each potential IT by integrating the Vienna Package ^14,18^ with default parameters into ITMiner. Specifically, ITMiner identified the most likely 2D folding structure of each IT (the structure with lowest change in free energy, Δ*G*) and obtained the sequence of the stem, loop, and poly-U tract (Supplementary Data 1 and 2). Stem length count was based on the total number of nucleotides on both sides of the stem. To deal with the issue of overlapping sequences being identified as different ITs, if the sequences of two ITs overlapped >90% and contained poly-U tracts identified through ITMiner, we considered them to be the same IT acting in both directions (strands), bringing the final candidate list in *E. coli* to 644. We validated this list by comparing it with two experimental datasets: Conway *et al*. 2014 and RegulonDB (accessed November 2021).

Finally, we used the existing annotations in the queried genomes (NC_000913 for *E. coli* and NZ_CP012347.1 for *S. pullorum*) to identify the genomic context of each IT – the identity and distance to upstream and downstream genes using the PyBED tools package in Python. We compared the differences in distances between upstream and downstream genes using a two-tailed paired t-test. This, and all subsequent statistical tests, were done in R version 4.2.3.

### Measuring IT efficiency

To measure the efficiency of ITs, we developed a synthetic construct that was inserted into a low copy number plasmid (2–3 copies) with the SC101* origin ^20^. The construct (Fig.3A) consisted of: an inducible *P_LAC_* promoter, which was always fully induced with 1mM IPTG; a *cfp* gene; RNase3 sites; an IT sequence surrounded by unique restriction sites; and a *yfp* gene. The inclusion of RNase3 sites surrounding the IT sequence resulted in the removal of the IT sequence from mRNA and the creation of separate transcripts, one for *cfp* and one for *yfp*. This approach ensured that translation was not affected by IT sequences. We selected 130 ITs from the list of 644 identified in the *E. coli* K12 MG1655 genome. They were selected randomly except we did not select any bi-directional ITs, as the underlying mechanisms of their function might be different to ITs that act on only one strand. We synthesized them as individual oligonucleotides (Sigma Aldrich) and cloned them into the construct using the unique restriction sites BamHI and SalI. Each cloned plasmid was independently electroporated into electrocompetent *E. coli* MG1655. Then, successful cloning was verified through Sanger sequencing.

We did not succeed in cloning one of the chosen ITs, resulting in 129 *E. coli* ITs inserted into our experimental system. To measure their efficiency, three biological replicates of each mutant were grown overnight in 96-well plates, at 37°C shaking at 650 rpm in a microplate shaker in M9 medium, supplemented with 0.2% (w/v) glucose, 0.2% (w/v) casein amino acids, 25 μg/mL kanamycin, and 1 mM IPTG. Overnight cultures were diluted 1:1000 into black 96-well plates and grown until optical density at 600 nm (OD_600_) was approximately 0.05. Then, they were inserted in a Biotek Synergy H1 platereader, set at 37°C with orbital shaking (282 cpm) to measure the following: OD_600_, CFP fluorescence (433 nm excitation wavelength, 475 nm emission), and YFP fluorescence (515 nm excitation, 543 nm emission) every 15 min for 3 h. Each plate included one blank, one negative control (containing a backbone plasmid with no fluorescence genes) and one positive control (containing the construct without any IT inserted). All replicate measurements were randomized among different plates. For each replicate, we selected a single time point that was as close as possible to OD_600_ of 0.25 to ensure that all measurements were in as similar of growth stage as possible.

To calculate the efficiency of an IT, we used the following formula:

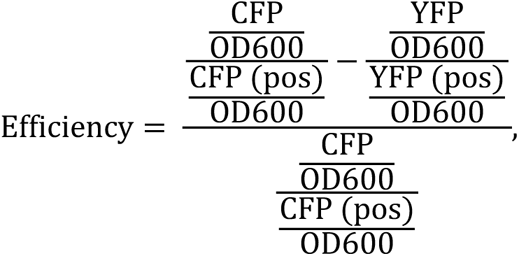

where CFP refers to the measured level of cyan fluorescence; YFP to the measured level of yellow fluorescence; (pos) to the measured levels in the respective positive controls; and OD600, to the measured optical density (600nm) of each sample. All values were first blank corrected. Based on this formula, efficiency values can range from zero to one, with zero being the lowest efficiency possible (no termination) and one, the highest (no polymerase read-through). They can also be negative when the sequence thought to be an IT acts as a promoter.

The efficiency of each IT was compared to a no-efficiency null hypothesis (efficiency of 0) by an FDR-corrected, one sample t-test, with P < 0.05 marking significant efficiency. To examine the effect of distance between and IT and its closest upstream or downstream gene, we conducted linear regressions with individual IT efficiency and distance as response variables.

The same protocol was used to measure the efficiency of 45 *S. pullorum* ITs, as well as 45 *E. coli* ITs in *S. pullorum*. The 45 *S. pullorum* ITs were selected semi-randomly, with the only conditions being that they have an identifiable ortholog in *E. coli* (see below) and that the two orthologs are not identical (*i.e.,* they must contain at least one difference in sequence). The comparison of efficiencies between orthologous ITs, or between the same IT measured in two hosts, was done using FDR-corrected, two sample t-tests.

### Conservation of ITs between E. coli and S. pullorum

We set out to create a comparison between two closely related species, *E. coli* and *S. enterica.* However, we originally obtained the genome and the IT list using ITMiner of *S. enterica* subsp. *pullorum* instead of serovar *Typhimurium* by mistake. We only realized this error once we have already measured the efficiency of all 45 *S. pullorum* ITs. Since the genomes are similar, we decided to go ahead with the data we already had on *S. pullorum*.

In order to find all homologs of *E. coli and S. pullorum* ITs in both genomes, we performed a comprehensive BLAST search: i) *E. coli* ITs against *E. coli* genome; ii) *E. coli* ITs against *S. pullorum* genome; iii) *S. pullorum* ITs against *E. coli* genome; and iv) *S. pullorum* ITs against *S. pullorum* genome. We used the following BLAST parameters to minimize false negatives at this stage: word_size = 4, E value >= 0.001. While most ITs had one or zero blast hits, some of them like Cha381 and Nur16 had many. The same procedure was applied to identify homologs and bidirectional best hits (BBHs) for protein-coding genes. The raw BLAST results are available in Supplementary Data 4.

Our main purpose was to find the best candidate for orthologous IT sequence in the non-native genome for each IT. To achieve this, we first applied filters to exclude duplicates and unrelated sequences (see below). Secondly, we searched for BBHs ^25^ for each *E.coli* IT among *S. pullorum* ITs and *vice versa*. Only one-to-one BBHs were considered orthologs. The following filters were applied: i) if two BLAST hits with similar score were mapped to opposite DNA strands and overlapped with each other and with an annotated IT, we considered only the hit in the same strand as the annotated IT; ii) if multiple BLAST hits were obtained, only the hit with the best score was taken into account. This method identified 106 orthologous pairs, or IT BBH pairs.

To find orthologous genes, we applied a similar procedure using the BBH approach with only the best BLAST hits taken into account. As a result, we found 3001 orthologous gene pairs (also referred to as gene BBHs) between *E. coli* and *S. pullorum*.

The all against all alignments of the 1,180 high-confidence IT sequences (644 from *E. coli*; 536 from *S. enterica*) was performed with the exact Smith-Waterman algorithm, using the ssearch36 binary version 36.3.8i from the fasta36 package ^34^. We visualized the results of the 1,392,400 pairwise alignments as a scatterplot −log10(default E-value) vs harmonic mean (sequence identity to query and query coverage) (Supplementary Fig.1). For stringency purposes, the harmonic mean was preferred over the arithmetic mean because it is skewed towards the lowest of the two values. The coordinates of BBHs identified with BLAST served as the basis to define the thresholds of −log10(E-value) >= 3 (corresponding to E-value <= 0.001) and harmonic mean (sequence identity, minimal coverage) >= 78% to justify an edge between two ITs in the sequence similarity network (SSN) (Supplementary Fig.1). The SSN was visualized and edited with Cytoscape version 3.9.1 ^35^, and analyzed with the R igraph package.

To create a phylogenetic tree of the homologous sequences of the two ITs with many hits in the non-native genome (Cha381 and Nur16), we first found all those hits using BLAST (as described above) and aligned them with Muscle with default parameters and visualized with AliView v 1.25 ^36^. The tree was constructed with MEGA X 10.1.8 ^37^ using the Neighbor Joining algorithm with default parameters (nucleotide, not protein-coding sequences, 1000 bootstrap iterations). The tree was visualized with FigTree v 1.4.4. Branches with small bootstrap values (< 0.1) were collapsed where possible.

## Acknowledgments

The authors would like to thank Nurdan Erdem and Tatjana Petrov for their bioinformatics support, and Nick Barton, Jonathan Bollback, Rok Grah, and Claudia Igler for the useful discussions. This work was funded by the Sir Henry Dale Fellowship jointly funded by the Wellcome Trust and the Royal Society (Grant Number 216779/Z/19/Z) to ML and FWF grant # I 3901-B32 to CCG.

## Author contributions

AN-S and ML conceived the study. CB developed ITMiner and conducted SSN analysis. SD carried out all other bioinformatics and genomics analyses with the support of RAFR. DT-A constructed experimental plasmids with the support of CCC and DAP. DT-A collected the experimental data and analyzed it together with ZK, AN-S, and ML. ML and SD wrote the first draft of the manuscript and revised it together with AN-S and CCG. All authors approved the final version of this manuscript.

## Supplementary Data

All supplementary data presented in the manuscript will be made available upon request.

