## Supplementary Table 1 and Figure 1 for "Evolution of intrinsic transcriptional terminators in their genomic context"

**Supplementary Table 1.** Amino acid sequence identity score between key molecules involved in transcriptional termination in *E. coli* and *S. pullorum*.

| **Molecule** | **Identity score (%)** |
|---|---|
| RNA polymerase subunit α | 99.70 |
| RNA polymerase subunit β | 99.85 |
| RNA polymerase subunit β' | 99.86 |
| termination factor NusG | 100.00 |
| Rho | 100.00 |
| antiterminator BglG | 98.74 |
| antiterminator NusB | 99.28 |
| sigma24 (sigmaE) | 100.00 |
| sigma28 (sigmaF) | 99.58 |
| sigmaS | 100.00 |
| sigma32 (sigmaH) | 100.00 |
| sigma54 | 99.79 |
| sigma70 (sigmaD) | 100.00 |

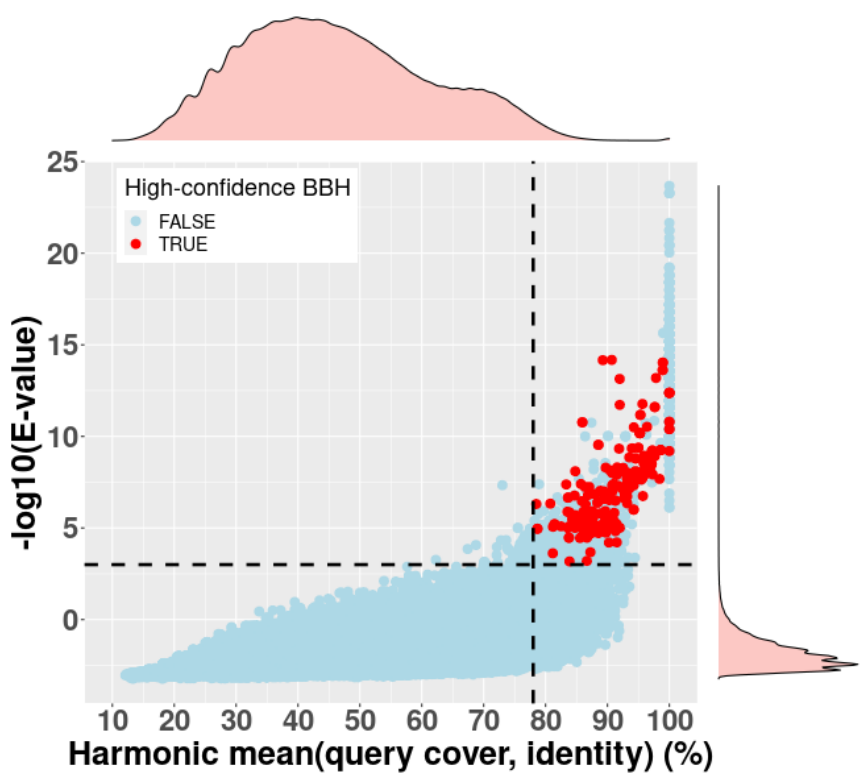


**Supplementary Figure 1.** All 1,392,400 pairwise Smith-Waterman All vs All alignments of the 1,180 high-confidence IT sequences (644 from *E. coli*; 536 from *S. pullorum*). Red dots correspond to best bidirectional hits (BBHs) between *E. coli* and *S. pullorum* identified with the BLAST method*,* serving as baselines to define the cutoffs of the IT Sequence Similarity Network (default E-value <= 0.001 and harmonic mean (sequence identity, minimal coverage) >= 78%).
